# *Luteibacter sahnii* sp. nov., a novel yellow-pigmented probiotic bacterium from rice seed microbiome

**DOI:** 10.1101/2023.09.28.560004

**Authors:** Gagandeep Jaiswal, Rekha Rana, Praveen Kumar Nayak, Rekha Chouhan, Sumit G. Gandhi, Hitendra K. Patel, Prabhu B. Patil

## Abstract

The genus *Luteibacter*, a member of the family *Rhodanobacteraceae*, encompasses Gram-negative bacteria found in diverse environments. In the present study, four yellow-pigmented bacterial isolates designated as PPL193^T^, PPL201, PPL552, and PPL554 were obtained and identified as Gram-negative, rod-shaped, and motile bacteria. Biochemical characterization and examination of the 16S rRNA gene sequence, derived from the genomic sequence, identified it as belonging to the genus *Luteibacter*. The isolates are closely related to *Luteibacter yeojuensis* R2A16-10^T^, forming a distinct monophyletic lineage with *L. aegosomatis* KCTC 92392^T^ and *L. anthropi* CCUG25036^T^. The calculated values for pairwise ortho Average Nucleotide Identity and digital DNA-DNA hybridization in comparison to previously reported *Luteibacter* species fell below the established thresholds for species delineation. As this novel species was isolated from rice seeds as a potential *Xanthomonas* due to its distinctive yellow-colored colonies, we sought to identify the presence of xanthomonadin pigment in this species. Intriguingly, our findings revealed the presence of the typical peak corresponding to xanthomonadin in the UV spectra, confirming its presence in this novel species and adaptation to plant habitat. Furthermore, the detailed genomic investigation also uncovered the genomic locus corresponding to xanthomonadin biosynthetic gene cluster, further suggesting that members of this novel species are co-habitants of plant pathogenic and plant probiotic *Xanthomonas* group of phytobacteria within rice seeds. Apart from protease production, the species was found to produce Indole-3-Acetic Acid (IAA) in higher quantities and was also able to protect plants from *Xanthomonas oryzae* pv. *oryzae*, a major pathogen of rice indicating its probiotic nature. Genome scanning revealed the presence of genomic region(s) encoding loci for biosynthesis of anti-microbial peptides and other metabolites with probiotic properties, further confirming its probiotic properties. This study highlights the importance of using a combination of phenotypic and genotypic methods for bacterial identification and expands our knowledge of the diversity and distribution of diverse bacteria associated with rice seeds and their microbiome. *Luteibacter sahnii* sp. nov. is proposed as a novel species of the genus *Luteibacter* with PPL193^T^=MTCC 13290^T^=ICMP 24807^T^=CFBP 9144^T^ as its type strain and PPL201, PPL552, and PPL554 as other constituent members.

## Introduction

*Luteibacter* is classified within the *Rhodanobacteraceae* family, positioned in the γ[subclass of Proteobacteria. This taxonomic group encompasses a diverse array of Gram-negative bacteria thriving in various unique habitats (Joe et al., 2023). The family *Rhodanobacteraceae* is closely related to the family *Lysobacteraceae* (*Xanthomonadaceae*). The genus *Luteibacter* is characterized as Gram[negative, rod-shaped, motile, aerobic bacteria with yellow-colored colonies (Friedrich et al., 2023a). Based on the very first species, *Luteibacter rhizovicinus* DSM 16549^T^, Johansen et al. first proposed the genus *Luteibacter* (Johansen et al., 2005). As of the time of writing, it consists of nine species: *L. rhizovicinus (Johansen et al., 2005)*, *L. yeojuensis (Kampfer et al., 2009)*, *L. anthropi (Kampfer et al., 2009)*, *L. jiangsuensis (Wang et al., 2011)*, *L. pinisoli (Akter and Huq, 2018)*, *L. flocculans (Friedrich et al., 2023a)*, *L. aegosomatis (Joe et al., 2023)*, *L. aegosomaticola (Joe et al., 2023)*, and *L. aegosomatissinici (Joe et al., 2023)*, of which three have been validly published (LPSN, http://www. bacterio.net) (Parte et al., 2020). Various *Luteibacter* species have been found in human blood, pond water, rhizospheric soil, greenhouse soil, and ecosystems linked with plants. *Luteibacter rhizovicinus* MIMR1 have even been uncovered to have the ability to facilitate plant growth (Guglielmetti et al., 2013). Members of this genus have DNA G+C content of 64.2–67.5 mol%. They are documented to exhibit positive results for oxidase and catalase tests and negative for urease tests (Akter and Huq, 2018).

*Xanthomonas* is an important plant-associated bacterium with a pathogenic and non-pathogenic lifestyle (Bansal et al., 2018). The non-pathogenic members are hypothesized to be diverse and complex members of the rice microbiome with probiotic properties. In a recent study, we reported that two non-pathogenic *Xanthomonas* (NPX) species can protect rice from a bacterial blight pathogen (Bansal et al., 2021; Rana et al., 2022; Rekha et al., 2023). Members of this genus are yellow, and the genus is named after the Greek word ‘Xantho,’ which means yellow. Considering the importance of *Xanthomonas*, we were interested in isolating new diverse strains and species of *Xanthomonas* from rice plants. During our screening for yellow-colored bacteria from healthy rice seeds, we isolated four yellow-pigmented strains. However, a phylogenetic investigation utilizing the sequence of 16S rRNA confirmed the strain’s affiliation with the *Luteibacter* genus. Interestingly, it is also named after its yellow coloras ‘Leuti’ in Latin means yellow, and phylo-taxonogenomic investigation revealed it to be a novel species. Further genome scanning unveiled the presence of a biosynthetic gene cluster related to xanthomonadin biosynthesis, and subsequent qualitative analysis confirmed the presence of an absorbance peak corresponding to xanthomonadin pigment.

Genome scanning also revealed the presence of genomic regions for the biosynthesis of antimicrobial compounds that include a lanthi-peptide and non-ribosomal peptide synthetase biosynthetic gene cluster (BGC). The strains were non-pathogenic and were able to protect the rice plants from *X. oryzae* pv. *oryzae* (Xoo), the leaf blight pathogen of rice. We could also find that the species was able to produce Indole-3-acetic acid (IAA) and also had protease activity, indicating its plant probiotic properties, which was further supported by genome scanning that revealed the presence of several additional genomic regions known to encode metabolites for plant probiotic properties. To our knowledge, this is the first report of a *Luteibacter* species isolated from rice seed, positing it as an important ecological and taxonomic counterpart of the *Xanthomonas* group of phytobacteria. It is pertinent to note that *Xanthomonas* and *Luteibacter* belong to *Lysobacteraceae* and *Rhodanobacteraceae* families, respectively. Both genera are closely related and fall within the order *Lysobacterales* (*Xanthomonadales*) (LPSN, https://lpsn.dsmz.de) (Parte et al., 2020). The genomic resource, along with comparative genomic and evolutionary findings, will be invaluable in future studies to understand rice seed microbiome in general and the *Xanthomonas* group of bacteria in particular.

## Materials and Methods

### Bacterial Strain Isolation

During the rice harvest season, rice seeds were collected from the rice fields in Ropar (30.9664°N, 76.5331°E) and Hyderabad (17.3871°N, 78.4917° E), India for bacterial isolation. To recover bacterial colonies, a well-established procedure described in the literature was followed, which involved crushing and spreading the seeds on four different media, namely Peptone Sucrose Agar (PSA), Nutrient Agar (NA), Tryptone Soya Agar (TSA), and Glucose Yeast Extract Calcium Carbonate Agar (GYCA) (Cottyn et al., 2001; Midha et al., 2016). The colonies on the media were carefully observed for their morphological characteristics, and those exhibiting a mucoid yellow color were selected for further analysis. To obtain pure cultures, single colonies were streaked onto the NA medium. The resultant pure cultures were then stored in 15% glycerol at −80°C for long-term preservation and subsequent experiments.

### Bacterial Ultrastructure Visualization by Transmission Electron Microscopy

To visualize the morphology of the isolates, transmission electron microscopy (TEM) was employed. To begin with, overnight cultures of the bacterium were grown on Nutrient Agar (NA) at a temperature of 28°C. Fresh bacterial colonies were then resuspended in Nutrient Broth (NB) and centrifuged at 2000 revolutions per minute (rpm) for a duration of 10 minutes. The resulting cell pellet underwent two washes with 1X phosphate-buffered saline (PBS) sourced from Gibco, USA, and then suspended in 50 µL of the same solution. Subsequently, by placing 10-20 µL of the bacterial suspension on a 300-mesh, carbon-coated copper grid (Nisshin EM) and allowed to rest for a period of 15 minutes. Next, the grid was subjected to negative staining using a 2% solution of phosphotungstic acid for a duration of 30 seconds, dried, and imaged with a JEOL JEM 2100 (Tokyo, Japan) operating at a voltage of 200 kV. This method enabled the visualization of the morphology of our isolates in great detail, providing insights into the structural characteristics of these bacteria.

### Phenotypic and Biochemical Characterization of *Luteibacter* Isolates

To examine the phenotypic characteristics, the *Luteibacter* isolates were grown in NA medium at a temperature of 28°C. The KOH test was conducted using the procedure described by Suslow et al. (Suslow et al., 1982a). The BIOLOG GN3 microplate system, developed by Biolog Inc (United States), was employed following protocol A provided by the manufacturer to conduct biochemical characterization of novel strains. This characterization encompassed the evaluation of carbon utilization, acid production, and a range of enzymatic activities. The microplates were positioned in an incubator at a controlled temperature of 28°C, and readings were intermittently taken at 24-hour intervals using the MicroStation 2 reader. Subsequently, the obtained results were subjected to analysis utilizing the MicroLog 3/5.2.01 software. The tests were repeated three times, with only the ± values being considered trustworthy. Notably, the outcomes at the 96-hour mark exhibited greater consistency and were, therefore, selected for further analysis.

### Whole-Genome Sequencing and Analysis

Genomic DNA from PPL193^T^, PPL201, PPL552, and PPL554 of bacterial cultures grown overnight in NB media overnight was extracted using the QuickDNA™ Fungal/Bacterial Miniprep Kit sourced from Zymo Research, United States. The quantity of the genomic DNA was assessed via Nanodrop 1000 from Thermo Fisher Scientific’s Invitrogen. Subsequently, the genomic DNA was sequenced using the Illumina NovaSeq (MedGenome, Hyderabad, India). The quality of the raw data was assessed using FastQC version 0.11.9 (https://qubeshub.org/resources/fastqc). The paired-end reads obtained from sequencing were subsequently assembled utilizing SPAdes version 3.15.5 (Prjibelski et al., 2020). To evaluate genome quality, we used the Quast version 5.2.0 tool (Gurevich et al., 2013) and CheckM version v1.2.2 (Parks et al., 2015). The annotation of draft genomes was conducted through utilization of the Prokaryotic Genome Annotation Pipeline (PGAP) available at NCBI (https://www.ncbi. nlm.nih.gov/genome/annotation_prok/).

### Phylo-taxonogenomic Analysis

To determine the phylogenetic relationship of the novel isolates PPL193^T^, PPL201, PPL552, and PPL554,a complete 16S rRNA and core gene mid-rooted phylogeny approach was used, which allowed to compare the sequences of these isolates with the type strains from the genus to gain insights into their evolutionary history and relatedness.

In order to analyze the 16S rRNA gene sequences of the bacterial isolates, the barrnap v0.9 tool was utilized, which was sourced from http://github.com/tseemann/barrnap. This software was used to retrieve complete gene sequences from both our isolates, type strains of other *Luteibacter* species, as well as *Fulvimonas soli* LMG19981, which was used as an outgroup. The sequences were cross-referenced with the EZBioCloud database to facilitate their identification (Yoon et al., 2017). The percentage identity of the isolates’ 16S rRNA gene sequence with other *Luteibacter* species was determined using the BLASTn program. The 16S rRNA gene sequences were then aligned and trimmed using the Muscle program (Edgar, 2004) and trimAl (Capella-Gutiérrez et al., 2009), respectively, using their default parameters to eliminate gaps and create a phylogenetic tree. The resulting neighbor-joining tree was constructed using the Jukes-Cantor model of nucleotide replacement, gamma-distributed rates among sites, and 1000 bootstrap repetitions in MEGA-X version v11 (Kumar et al., 2018). The phylogenetic tree was subsequently annotated using iTOL v6 (Letunic and Bork, 2021).

To generate the core gene phylogeny, the whole-genome sequences of all the type strains from the genus *Luteibacter* were obtained from the NCBI GenBank database. Roary v3.12.0 (Page et al., 2015), which allows for the fast and accurate identification of orthologous gene clusters in large-scale bacterial datasets, was employed to create a concatenated alignment of the core genes based on the Prokkav1.14.6 (Seemann, 2014) annotated files. Subsequently, PhyML v3.3.20211231 (Guindon et al., 2010) was used to generate a maximum likelihood phylogeny based on this alignment file. For this, the General Time Reversible (GTR) nucleotide substitution model, which is a commonly used model for phylogenetic analysis of DNA sequences, was used. Additionally, we used 1000 bootstrap replicates to ensure the robustness of the phylogenetic tree. Finally, we used iTOL v6 (Letunic and Bork, 2021) to visualize and annotate the phylogenetic tree.

The orthoANI values were computed through the oANI method using USEARCH version 11.0.667 (Lee et al., 2016), while the dDDH values were determined utilizing formula 2 from the Genome-to-Genome Distance Calculator version 3.0 (Meier-Kolthoff et al., 2022).

### Isolation and characterization of xanthomonadin pigment and biosynthetic locus

PPL193^T^, PPL201, PPL552, PPL554, *X. oryzae, X. citri, X. sontii,* and *X. viticola* were grown in NA medium at a temperature of 28°C. The bacterial cells were scrapped off from the surface of NA plates and washed with PBS for 2-3 times. The cell pellet was then collected by centrifugation. Pigments were extracted by incubating the cell pellet with 1.5 mL of methanol at 60°C for 5 minutes in shaking conditions. The cells were pelleted by centrifugation at 8000 × g for 15 minutes. The clear supernatant was the methanolic extract, which contained xanthomonadin pigment. The spectra were observed using a UV/Visible spectrophotometer (200 to 800 nm), and the quantity of pigment generated was plotted as absorbance (OD at 445 nm) (Poplawsky and Chun, 1997; Soudi et al., 2011).

To identify the xanthomonadin biosynthetic gene cluster in PPL193^T^ and other previously reported *Luteibacter* species, we began by obtaining gene amino acid sequences from previously known xanthomonadin BGC (from *X. oryzae* and *X. campestris*) on NCBI through locus tags. These sequences were then used as a query to search for their homologs using tBLASTn v2.9.0+. The thresholds for percent identity and query length coverage were kept at values greater than 40% to ensure the accuracy of the process. Clinker v0.0.23 178 (Gilchrist and Chooi, 2021) was utilized to generate comparative gene clusters using BLASTp.

### Plant growth promotion studies: Enzymatic and hormonal profiling of *Luteibacter* strain PPL193^T^

To assess the plant growth promotion capabilities of *Luteibacter* PPL193^T^, both protease and Indole-3-Acetic Acid (IAA) production were evaluated. Protease production was assessed qualitatively on a 1% Skimmed Milk Agar (SMA) medium. A colony of PPL193^T^ was spot-inoculated onto the SMA plate and incubated at 28°C for ten days. A clear zone around the colony indicated the presence of protease activity. The diameter of the clear zone was measured to quantify the protease activity. The production of IAA by PPL193^T^ was assessed using spectrophotometry. PPL193^T^ was inoculated into 50 mL of Nutrient Broth (NB) containing 0.2% L-tryptophan and then subjected to incubation at 28°C with agitation at 125 rpm for seven days. The culture broth was then centrifuged (10000 rpm), and to 500 µL of the supernatant, 1 mL of Salkowski reagent was added. After one hour of incubation, the development of a pink color indicated the presence of IAA. The optical density (OD) was measured at 530 nm to quantify IAA production. A standard curve of IAA was used to determine the concentration of IAA produced by PPL193^T^.

### *In planta* virulence and co-inoculation studies

To evaluate the pathogenicity of two distinct isolates among the four newly identified *Luteibacter* isolates, namely PPL193^T^ and PPL552, clip inoculation experiments were conducted on rice leaves. In brief, bacterial cultures were cultivated overnight in PS media (composed of 1% peptone and sucrose, pH 7.2), followed by centrifugation, washing with Milli-Q water, and adjustment to an OD_600nm_1. Clip inoculation was performed on leaves of rice plants aged between 60 to 80 days, using the rice cultivar Taichung Native 1 (TN1). As a positive control, the BXO1 strain of *X. oryzae* pv. *oryzae* (Xoo), a known rice pathogen, was employed, while Milli-Q served as the neutral/negative control. The lesion lengths were recorded 15 days post-inoculation, considering a minimum of 8 leaves. Average lesion lengths were depicted on a graph with standard deviations as error bars. Statistical significance was assessed using the Student’s t-test. This virulence assay was replicated three times. Furthermore, the novel strains were examined to check whether they provide protection against Xoo infection through co-inoculation experiments. Rice leaves from 60 to 80 days old TN1 plants were used for this purpose. Similar to the virulence assay, bacterial cultures were grown overnight in PS media, pelleted, and washed with Milli-Q water. The *Luteibacter* bacterial suspension, equivalent to an OD_600nm_ 1/mL, was adjusted to 0.5mL volume and then mixed with an equivalent volume (OD_600nm_ 1 in 0.5 mL) of BXO43 suspension immediately before clip inoculation. This ensured that both *Luteibacter* and Xoo were present in the final mixture, with an OD_600nm_ of 1, within a 1ml culture volume.For control groups, rice leaves were clip-inoculated using scissors immersed in either Milli-Q water or an OD_600nm_ of 1 BXO43 suspension. Lesion lengths were measured 15 days post-inoculation, considering a minimum of 8 leaves, and average lesion lengths were graphed with standard deviations as error bars. Similar to the virulence assay, the co-inoculation experiment was performed in triplicates.

### Genomic insights into secondary metabolite biosynthesis and IAA biosynthesis-related gene prediction

Genomes of PPL193^T^ and PPL552 were analyzed with antiSMASH bacterial version 7.0.1 (https://antismash.secondarymetabolites.org) with default settings to identify the presence of potential secondary metabolites or BGCs associated with more than 20 natural product classes such as ribosomally synthesized and post-translationally modified (RiPP-like) peptides, non-ribosomal peptide synthetase (NRPS) peptides, polyketide synthases (PKS) compounds, lanthipeptides, etc. (Blin et al., 2021).

In order to predict the presence of the indole-3-acetamide hydrolase (IaaH) gene within novel *Luteibacter* species, a tblastn (version 2.14.1+) analysis was conducted, employing a reference sequence obtained from the UniProt database (P06618). The genomic sequences of the four strains of the novel species served as subject databases for the tblastn search. Specific criteria, including an e-value threshold of ≤ 1e-5 and a minimum alignment length of ≥ 60%, to identify significant homologous sequences were applied.

### Pangenome Analysis

The pangenome analysis aimed to identify genes shared among *Luteibacter* species and unique to the novel isolate PPL193^T^ was conducted using Roary v3.13.0 (Page et al., 2015). GFF files generated by Prokka v1.14.6 (Seemann, 2014) were used as input for Roary with default parameters and a BLASTp identity cut-off of 90%. The functional annotation of unique genes was conducted to gain insights into the functional attributes of PPL193^T^ and its closely associated species *L. aegosomatis* KCTC 92392^T^ using eggNOG-mapper v2, which is an integral part of the eggNOG v5.0 database (Huerta-Cepas et al., 2019). The genes were categorized into clusters of orthologous groups (COG) categories, providing valuable information about their putative functions and potential biological roles.

## Results and Discussion

### Cellular, biochemical, and phenotypic characterization of yellow-pigmented bacterial isolates from rice seed

Four bacterial isolates, namely PPL193^T^, PPL201, PPL552, and PPL554, were obtained from healthy rice seeds. These isolates formed pale yellow mucoid textured spherical colonies after 24-48 hours of growth at 28°C on NA medium. The isolates were also found to be Gram-negative, as indicated by the thick, stringy, and long strand formation within the first 30 seconds of the KOH test (Suslow et al., 1982b). Morphologically, TEM data revealed that the cells exhibited straight rods with rounded ends and motility via a single polar flagellum (**Fig. 1**). The selection of the four strains, PPL193^T^, PPL552, PPL554, and PPL201, was based on their distinctive yellow-colored mucoid colonies, as well as their potential as non-pathogenic *Xanthomonas* strains found in rice seeds (Bansal et al., 2021; Rana et al., 2022). Subsequently, the pigments produced by these strains were isolated, and their spectral properties were compared with the reported spectra known for xanthomonadin (Rajagopal et al., 1997). *X. oryzae, X. citri, and X. sontii* were taken as positive controls as they possess the xanthomonadin biosynthetic gene cluster (BGC). Meanwhile, *X. viticola* that exhibit a frameshift mutation in a gene encoding phospho-transferase, leading to a truncated protein with 122 amino acids instead of the native 143 amino acids (deletion of four nucleotide bases at positions 348–351), consequently manifesting a non-pigmented colony phenotype was taken as the negative control to validate the absence of the xanthomonadin peak (Rana et al., 2023). Notably, we observed characteristic humps in the obtained spectra, which are consistent with those typically associated with xanthomonadin (**Fig. 2**).

**Fig. 1:**
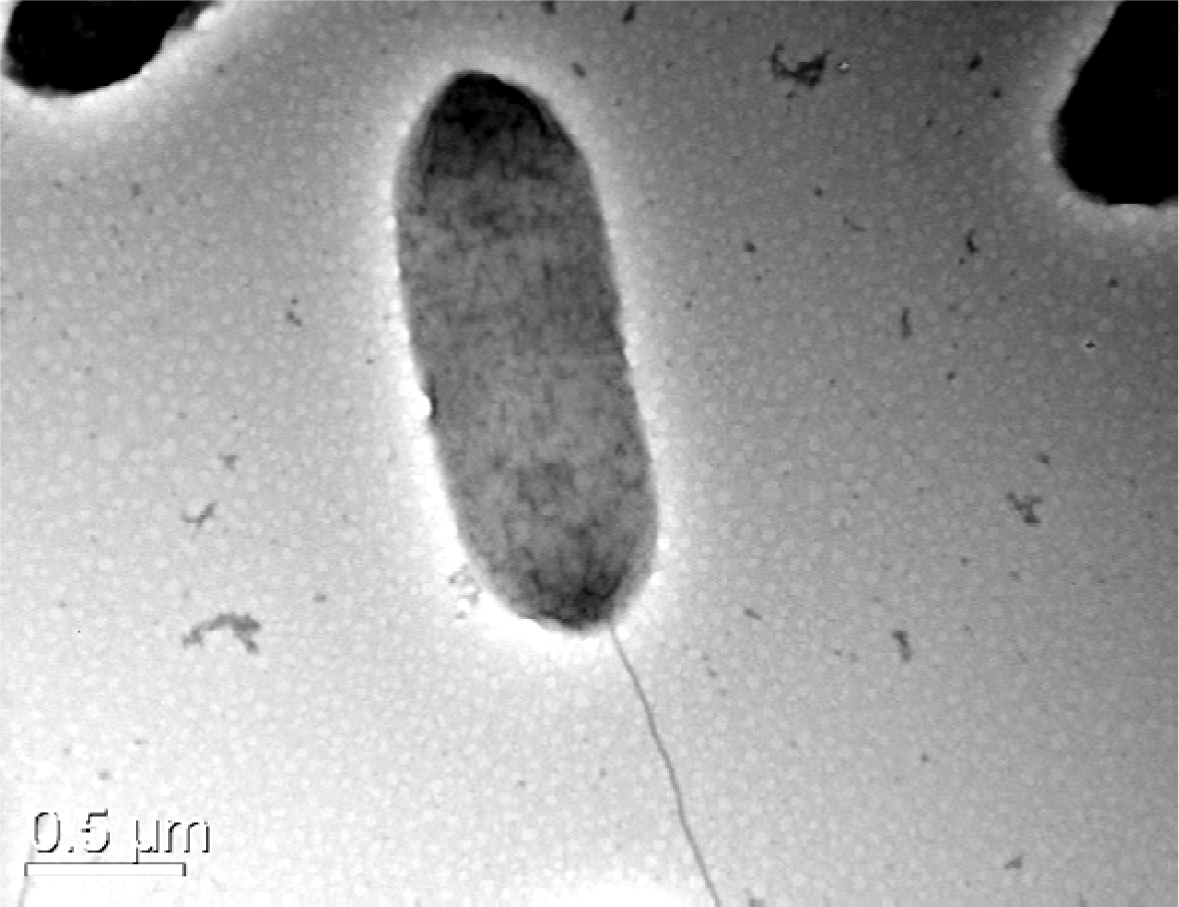
Transmission Electron Microscopy revealed the intricacies, showcasing the flagellated, rod-shaped morphotype of the *Luteibacter* PPL193^T^ isolate. This micrograph was captured after 24 hours of cultivation on nutrient agar (NA) at 28°C, followed by negative staining, unveiling exquisite surface details and cellular morphology.

**Fig. 2:**
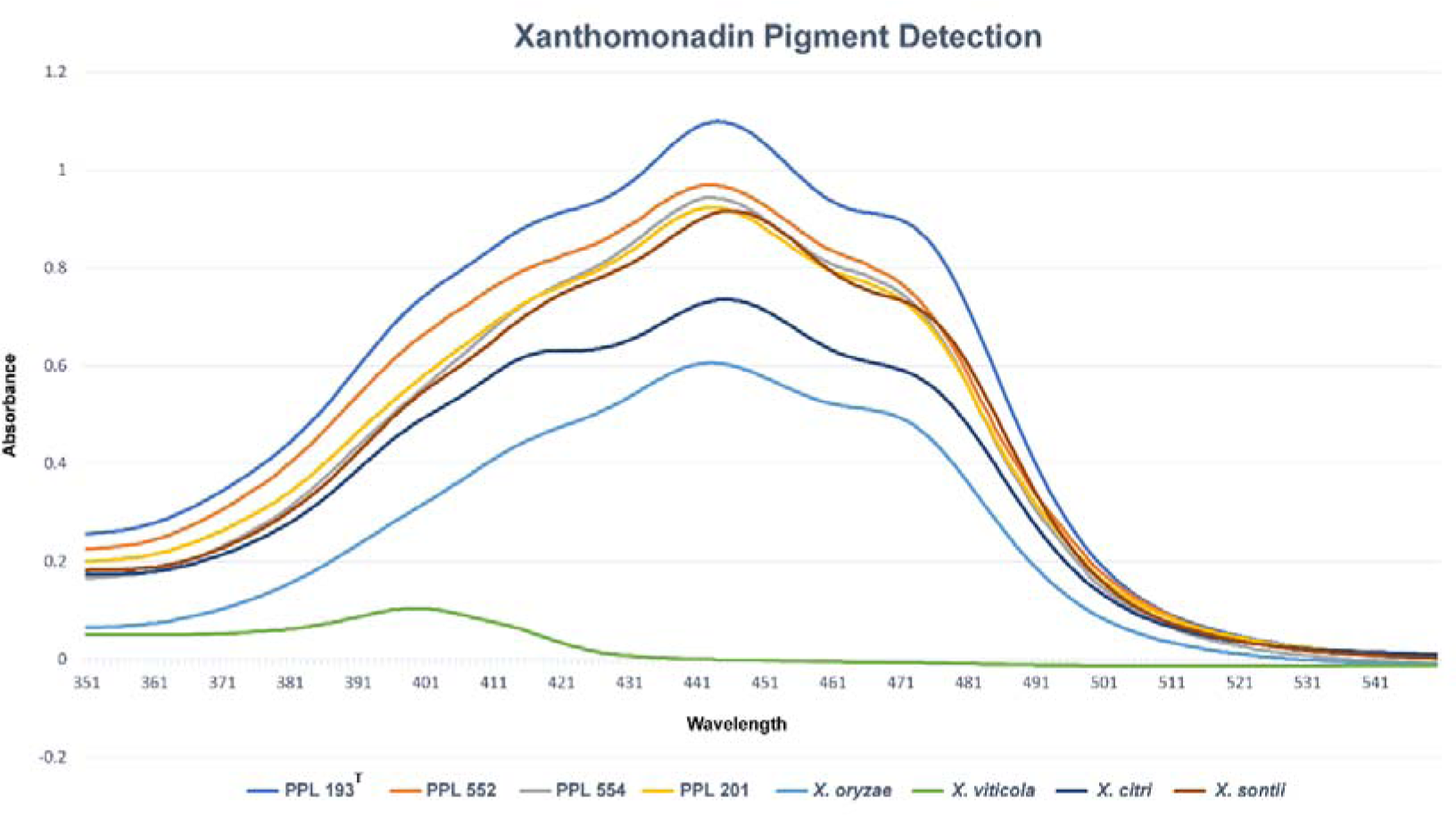
Detection of xanthomonadin pigment in novel *Luteibacter* strains (PPL193^T^, PPL201, PPL552, and PPL554).For validation, *X. oryzae, X. citri, and X. sontii* were employed as positive controls due to their possession of the xanthomonadin biosynthetic gene cluster (BGC), while *X. viticola*, lacking the xanthomonadin BGC due to frameshift mutation in one of the genes, was designated as a negative control.

In order to obtain a comprehensive understanding of the biochemical characteristics of the newly isolated strain PPL193^T^, a series of biochemical tests were conducted using the OMNILOG GEN III system (BIOLOG). The results of these tests, as shown in **Table 1**, reveal that the novel strain has a unique metabolic profile that is distinct from other *Luteibacter* species. Specifically, PPL193^T^ was found to be capable of utilizing a variety of carbohydrates, including D-trehalose, D-cellobiose, gentiobiose, sucrose, D-turanose, α-D-lactose, N-Acetyl-D-Glucosamine, α-D-glucose, D-mannose, L-fucose, D-maltose, stachyose, D-melibiose, and β-Methyl-D-Glucoside, among others. The strain was found to be resistant to several antibiotics, such as nalidixic acid, lincomycin, rifamycin SV, and aztreonam. In addition to carbohydrates, PPL193^T^ was also able to utilize a range of other substrates, including L-alanine, L-glutamic acid, L-serine, tetrazolium violet, tetrazolium blue, and L-lactic acid. Comparison of PPL193^T^ with other *Luteibacter* species (Akter and Huq, 2018; Friedrich et al., 2023a; Friedrich et al., 2023b; Joe et al., 2023; Johansen et al., 2005; Kampfer et al., 2009; Wang et al., 2011) revealed significant differences in the metabolism of certain compounds such as α-D-glucose, D-fucose, inosine, L-Malic Acid, α-keto-butyric acid, and propionic acid **(Table 1)**.

**Table 1:**
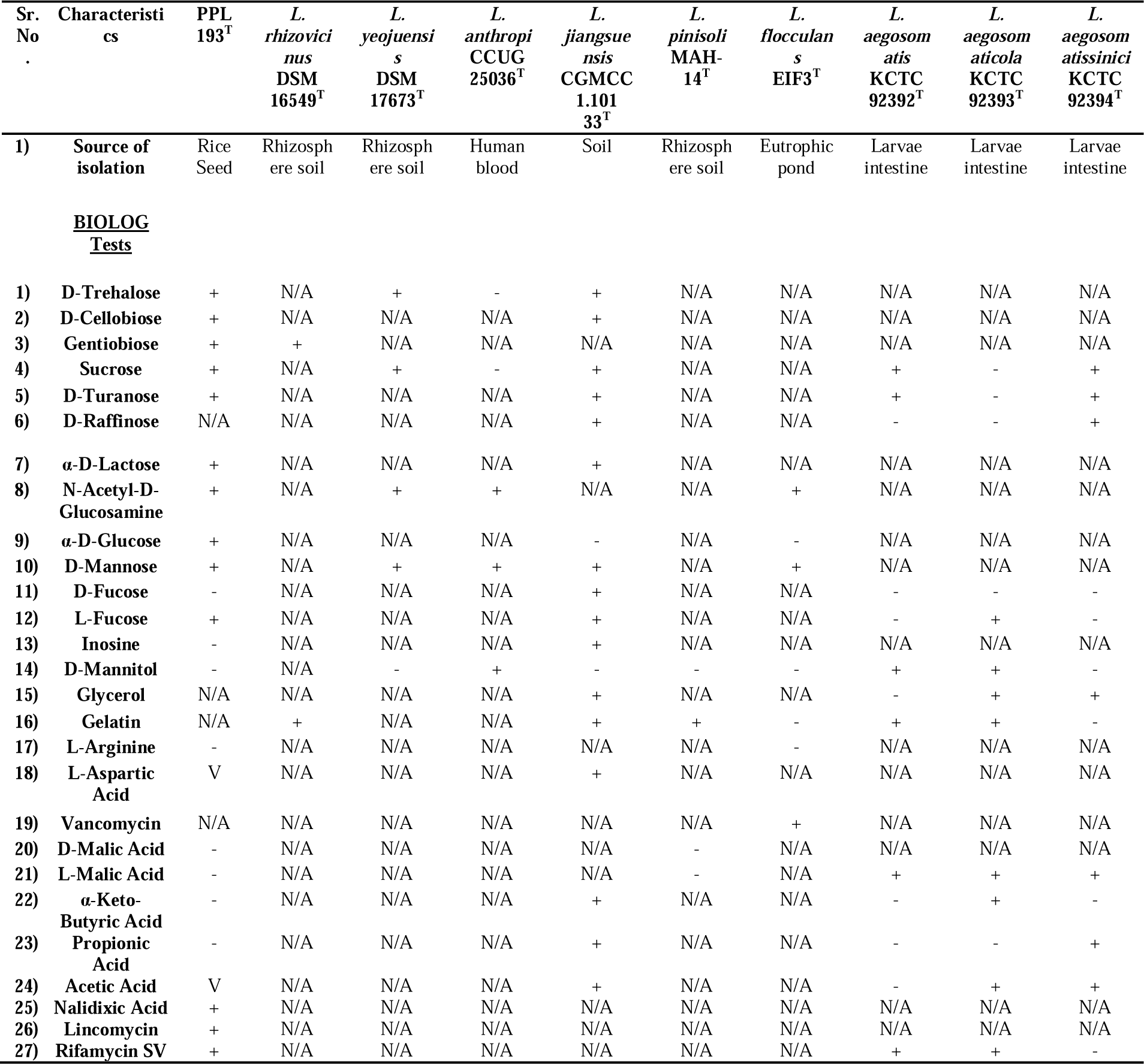
Distinguishing biochemical characteristics: Comparative analysis of PPL193^T^ and phylogenetically related *Luteibacter* species. Investigating key traits between 1) strain PPL193^T^ and Taxa: 2) *L. rhizovicinus* DSM 16549^T^, 3*) L. yeojuensis* DSM 17673^T^, 4) *L. antrophi* CCUG 25036^T^, 5) *L. jiangsuensis* CGMCC 1.10133^T^, 6) *L. pinisoli* MAH-14^T^, 7) *L. flocculans* EIF3^T^, 8) *L. aegosomatis* KCTC92392^T^, 9) *L. aegosomaticola* KCTC92393^T^, 10) *L. aegosomatissinici* KCTC92394^T^. Biochemical Traits: + (Positive), - (Negative), N/A (Data Not Available), V (Variable).

### Phylo-Taxonogenomic analysis of PPL193^T^, PPL201, PPL552, and PPL554

The genomic characteristics of PPL193^T^, PPL201, PPL552, and PPL554 were explored using genome-based taxonomic and phylogenetic analysis. The total genome size of these strains was found to be 4.21 Mbp, 4.24 Mbp, 4.17 Mbp, and 4.19 Mbp, respectively, with a GC content of approximately 67%. The genome sequences were assembled into 36, 49, 29, and 29 contigs, and the average/mean genome coverage and N50 values were 942x, 535x, 515x, and 530x and 11,34,670, 12,17,826, 3,27,166, and 10,39,592 bp, respectively (**Table 2)**.The complete 16S rRNA gene sequence of the isolates obtained from the whole genome sequencing revealed the highest identity to *Luteibacter yeojuensis* R2A16-10 in the EZBioCloud database (Yoon et al., 2017). The 16S rRNA gene sequences of the isolates exhibited complete similarity with one another and over 98% similarity with *Luteibacter* species within clade II (**Table. 3**). This clade includes well-known members such as *L. rhizovicinus*, *L. anthropi*, *L. pinisoli*, *L. yeojuensis*, *L. aegosomatis*, *L. aegosomaticola*, and *L. aegosomatissinici*. Furthermore, the phylogenetic tree based on 16S rRNA gene sequences of the isolates and other related *Luteibacter* species while taking *F. soli* LMG19981^T^ (Mergaert et al., 2002) as an outgroup, which revealed that the novel isolates formed a monophyletic clade among clade II *Luteibacter* species (**Fig. 3**).

**Fig. 3:**
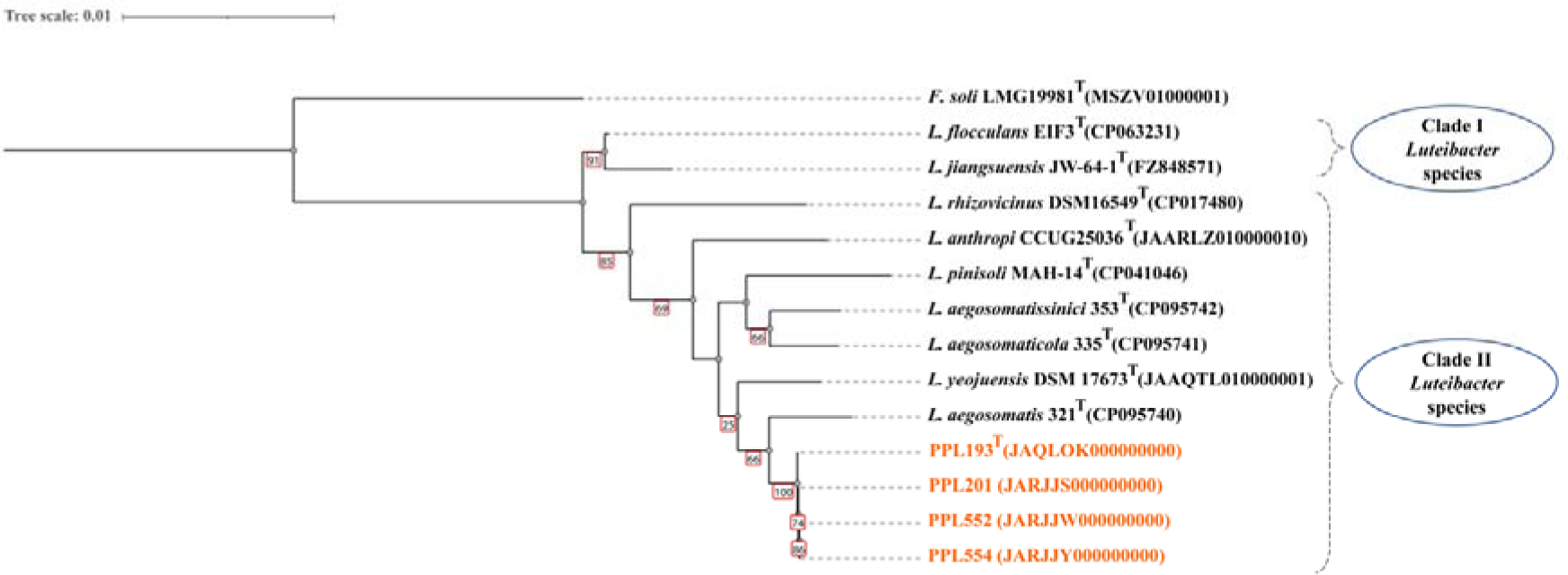
Neighbor-joining analysis of 16S rRNA gene sequences. The constructed phylogeny utilizing 16S rRNA gene sequences provides insights into the evolutionary connections among the novel isolates and other *Luteibacter* species. The NCBI GenBank database accession numbers, enclosed in brackets after the species names, authenticate the identification. PPL193^T^, PPL201, PPL552, and PPL554 are visually highlighted in vibrant orange color. The red-bordered rectangles at the nodes display robust bootstrap values, emphasizing confidence in the inferred relationships. Omitting values below 50%, the scale bar represents the number of nucleotide substitutions per site, providing a visual estimation of genetic diversity.

**Table 2.** Genome assembly and annotation metrics: Illuminating the genomic profiles of PPL193^T^, PPL201, PPL552, and PPL554.

| STRAINS | GENOME<br>SIZE | GC<br>CONTENT<br>(%) | GENOME<br>COVERAGE<br>(X) | CONTIGS | N50 (BP) | CDS | RNAS | PSEUDO-GENES | ACCESSION<br>NUMBER |
| --- | --- | --- | --- | --- | --- | --- | --- | --- | --- |
| PPL193<sup>T</sup> | 4211065 | 67.00 | 942 | 36 | 1134670 | 3,634 | 57 | 32 | JAQLOK000000000 |
| PPL201 | 4244686 | 66.95 | 535 | 49 | 1217826 | 3,677 | 59 | 36 | JARJJS000000000 |
| PPL552 | 4165127 | 67.10 | 515 | 29 | 327166 | 3,617 | 60 | 35 | JARJJW000000000 |
| PPL554 | 4189012 | 67.03 | 530 | 29 | 1039592 | 3,635 | 58 | 39 | JARJJY000000000 |

To establish the taxonomic position of PPL193^T^, PPL201, PPL552, and PPL554, the genomic indices were calculated with type strains of already reported species, including three validly published species and recently reported *Luteibacter* strains. The OrthoANI and dDDH values of these strains were found to be ≤83.8% and <30%, respectively, with other species of the genus *Luteibacter* (LPSN, https://lpsn.dsmz.de/genus/luteibacter) (Parte et al., 2020), which is below the species delineation cut-offs. The closest relatives of these strains were found to be *L. aegosomatis*, *L. anthropi*, and *L. yeojuensis*, with their orthoANI values ranging from 80% to 83.8%. The orthoANI values between the isolates themselves were in the range of 99.2-99.8%, and the dDDH values were in the range of 94-98.1% (**Table 3**) (Kim et al., 2014).

**Table 3:** Comparative analysis of genetic relatedness: Assessment of digital DNA-DNA Hybridization (dDDH), Average Nucleotide Identity (ANI), and 16S rRNA gene similarity. Exploration of the genomic proximity of PPL193^T^, PPL201, PPL552, and PPL554 isolates with representative strains from various *Luteibacter* species. Calculated values are expressed as percentages.

| Sr. No. | Strains | dDDH |  |  |  | ANI |  |  |  | 16S rRNA identity |  |  |  |
| --- | --- | --- | --- | --- | --- | --- | --- | --- | --- | --- | --- | --- | --- |
|  |  | PPL193<sup>T</sup> | PPL552 | PPL554 | PPL201 | PPL193<sup>T</sup> | PPL552 | PPL554 | PPL201 | PPL193<sup>T</sup> | PPL552 | PPL554 | PPL201 |
| 1. | PPL193<sup>T</sup> | 100 | 94.1 | 95.3 | 95.6 | 100 | 99.2283 | 99.4005 | 99.4097 | 100 | 100 | 100 | 100 |
| 2. | PPL552 | 94.1 | 100 | 94.2 | 94 | 99.2283 | 100 | 99.2756 | 99.2094 | 100 | 100 | 100 | 100 |
| 3. | PPL554 | 95.3 | 94.2 | 100 | 98.1 | 99.4005 | 99.2756 | 100 | 99.7184 | 100 | 100 | 100 | 100 |
| 4. | PPL201 | 95.6 | 94 | 98.1 | 100 | 99.4097 | 99.2094 | 99.7184 | 100 | 100 | 100 | 100 | 100 |
| 5. | *L. rhizovicinus* DSM 16549<sup>T</sup> | 22.5 | 22.5 | 22.6 | 22.5 | 79.1644 | 79.2405 | 79.1619 | 79.0808 | 98.45 | 98.36 | 98.45 | 98.45 |
| 6. | *L. anthropi* CCUG 25036<sup>T</sup> | 24.2 | 24.3 | 24.3 | 24.2 | 80.9156 | 80.9134 | 80.8281 | 80.8254 | 98.96 | 98.90 | 98.96 | 98.96 |
| 7. | *L. yeojensis* DSM 17673<sup>T</sup> | 23.5 | 23.6 | 23.5 | 23.5 | 80.2058 | 80.4492 | 80.0798 | 80.5027 | 99.35 | 99.32 | 99.35 | 99.35 |
| 8. | *L. jiangsuensis* CGMCC 1.10133<sup>T</sup> | 23.1 | 23.1 | 23.1 | 23.2 | 79.8489 | 79.7888 | 79.9017 | 79.7435 | 99.45 | 99.38 | 99.45 | 99.45 |
| 9. | *L. pinisoli* MAH-14<sup>T</sup> | 22.5 | 22.5 | 22.5 | 22.5 | 79.6631 | 79.9201 | 79.9164 | 79.801 | 98.77 | 98.77 | 98.77 | 98.77 |
| 10. | *L. flocculans* EIF3<sup>T</sup> | 22.5 | 22.5 | 22.5 | 22.6 | 79.4324 | 79.5608 | 79.3204 | 79.6194 | 99.03 | 98.97 | 99.03 | 99.03 |
| 11. | *L. aegosomatis*KCTC9 2392<sup>T</sup> | 27.4 | 27.4 | 27.4 | 27.4 | 83.7004 | 83.7493 | 83.802 | 83.8142 | 99.48 | 99.45 | 99.48 | 99.48 |
| 12. | *L. aegosomaticola*KC TC92393<sup>T</sup> | 22.3 | 22.3 | 22.4 | 22.4 | 79.1814 | 79.1432 | 78.977 | 79.2434 | 99.03 | 98.97 | 99.03 | 99.03 |
| 13. | *L. aegosomatissinici*K CTC 92394<sup>T</sup> | 22.3 | 22.3 | 22.3 | 22.3 | 78.9154 | 79.0039 | 78.8342 | 78.8884 | 99.03 | 98.97 | 99.03 | 99.03 |

### Investigating evolutionary relationships of *Luteibacter* through core gene-based phylogeny

In order to investigate the phylogenetic relationships of the novel *Luteibacter* isolates, a core gene-based analysis was performed. In the case of these isolates and the type strains of *Luteibacter* spp., the core gene content was found to consist of 651 genes. The next step was to use these core genes to construct a mid-rooted phylogenetic tree, which could reveal the evolutionary relationships between the strains. The resulting tree showed that the novel isolates belonged to clade II of the genus *Luteibacter*, which is consistent with their taxonomic classification (**Fig. 4**). However, the most exciting finding was that the novel strains formed a monophyletic sub-clade that was closely associated with *L. aegosomatis*, a recently reported species of *Luteibacter* from the intestine of insect larvae (Joe et al., 2023).

**Fig. 4:**
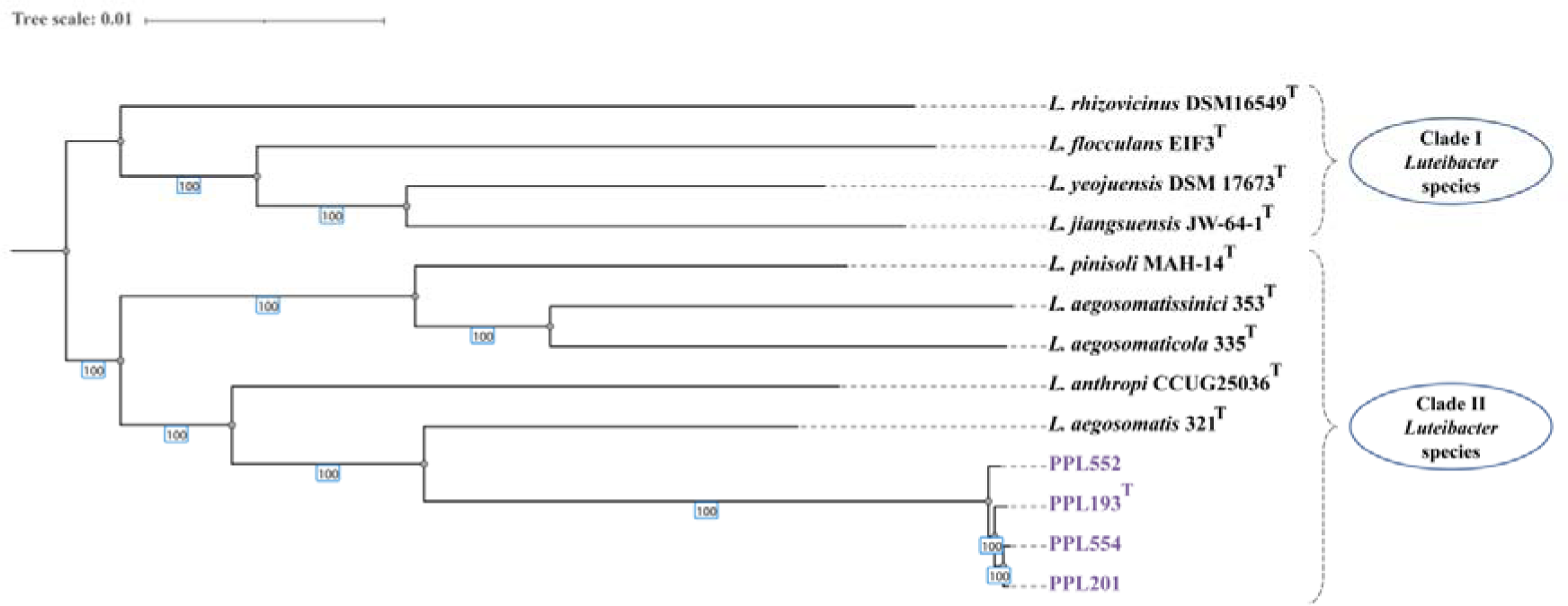
The maximum likelihood phylogenetic analysis of novel *Luteibacter* strains and related species was performed based on the core gene content of PPL193^T^, PPL201, PPL552, and PPL554, in conjunction with representative strains of other *Luteibacter* species. The scale bar within the figure illustrates the genetic diversity, showcasing the number of nucleotide substitutions per site. Notably, PPL193^T^, PPL201, PPL552, and PPL554 are visually highlighted in vibrant purple, drawing attention to their distinctiveness. Additionally, the bootstrap values, presented in distinctive blue-bordered rectangles, provide statistical support at the nodes, further reinforcing the robustness of the phylogenetic relationships depicted in this captivating visualization.

Overall, the core gene-based phylogenetic analysis presented in this study provides valuable insights into the taxonomic position and evolutionary relationships of the novel *Luteibacter* isolates. These findings have important implications for our understanding of bacterial diversity and evolution and could inform future studies aimed at exploring the genetic and ecological characteristics of this group of bacteria.

### Identification and comparative analysis of a putative xanthomonadin biosynthetic gene cluster

In our previous study, we highlighted the significance of the acquisition of the xanthomonadin biosynthetic gene cluster (BGC) in the evolution of *Xanthomonas* bacteria and their successful adaptation to plant hosts (Bansal et al., 2020). Recent reports, including our own investigations, have shed light on the existence of a hidden, diverse, and intricate world of non-pathogenic *Xanthomonas* (NPX) strains with probiotic properties, including the ability to protect plants against pathogen(s) (Bansal et al., 2021; Rana et al., 2022; Rekha et al., 2023; Triplett et al., 2015). We reported two NPX species from rice (Bansal et al., 2021; Rana et al., 2022) and expressed our interest in isolating and identifying additional NPX species from rice seeds. In this context, during culturomics of rice seed microbiome, our focus was on yellow-pigmented mucoid colonies exhibiting characteristic xanthomonadin spectral peaks, as they were potential candidates for NPX strains or species. However, the phylo-taxonogenomic analysis revealed that these novel strains belong to a new species from the genus *Luteibacter*. Along with isolation and characterization of pigment using UV/Visible spectroscopy, we also performed a comprehensive genome scan, leading to the identification of a potential xanthomonadin biosynthetic gene cluster (**Fig. 5**). The presence of xanthomonadin in our identified species indicates its importance as a prominent member of the rice microbiome, alongside NPX species. Notably, a recently published study on a novel species, *L. flocculans*, also reported the possibility of xanthomonadin biosynthesis in this genus, as evidenced by antiSMASH results (Friedrich et al., 2023a). In the comparative analysis of the xanthomonadin BGC (**Fig. 5**), we examined the composition of the cluster in our novel and other known *Luteibacter* species, as well as compared it to the xanthomonadin clusters of *Xanthomonas oryzae* and *Xanthomonas campestris*.

**Fig. 5:**
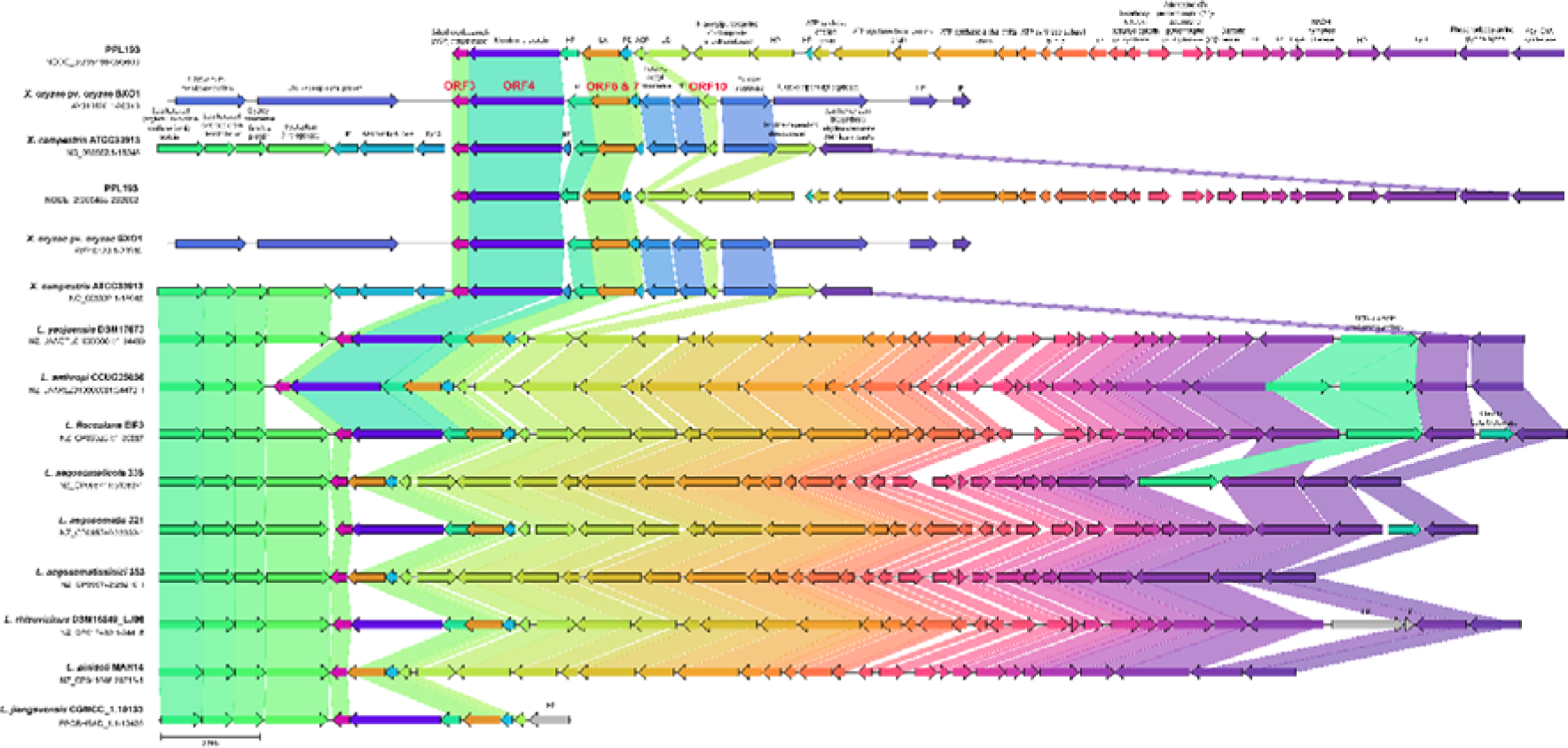
Comparative analysis of genomic loci involved in xanthomonadin biosynthetic gene clusters from PPL193^T^, *X. oryzae*, *X. campestris*, and other known *Luteibacter* species. Genes are depicted by arrows, and consistent colors are used to emphasize the presence of identical genes across multiple strains. Figure abbreviations include HP: Hypothetical Protein, PD: Putative Dehydratase, PH: Putative Halogenase, GF2 Protein: Glycosyl transferase Family 2 Protein, BKS Family Protein: Beta-Ketoacyl-[Acyl-Carrier-Protein] Synthase Family Protein, AP: Adenosine (5’)-Pentaphospho-(5’’)-Adenosine Pyrophosphohydrolase, ErpA: Iron-Sulfur Cluster Insertion Protein ErpA, LplT: Lysophospholipid Transporter.

Interestingly, we found that the xanthomonadin cluster of our novel *Luteibacter* species, as well as the previously characterized *Luteibacter* species, shares key similarities with the xanthomonadin clusters of *Xanthomonas oryzae* and *Xanthomonas campestris*. Specifically, we identified the presence of essential genes for xanthomonadin production and outer membrane localization of xanthomonadin, such as ORFs 3, 6, 7, 10, and ORF 4, which have been reported in a study elucidating the xanthomonadin cluster of *Xanthomonas oryzae* (Goel et al., 2002). Moreover, our analysis revealed the presence of novel genes within the BGC of our novel *Luteibacter* species, which were not previously reported in known xanthomonadin clusters from *Xanthomonas* species. These novel genes were found to encode proteins involved in various cellular processes, including N-acetylglucosamine-1-phosphate uridyltransferase, ATP synthase subunits, 6-carboxy-5,6,7,8-tetrahydropterin synthase, and other enzymes and transporters. These novel genes represent potential additions to the xanthomonadin biosynthetic pathway and may contribute to the unique properties and functionalities of the pigment produced by the novel *Luteibacter* species.

### Insights from plant growth promoting traits

*Luteibacter* strain PPL193^T^ exhibited a versatile enzymatic profile, showcasing both protease and IAA production capabilities. The protease production assay revealed the presence of a clear zone with a radius of 1.36 ± 0.057 cm around the colony on 1% Skimmed Milk Agar (SMA) medium, signifying its ability to produce proteases (**Fig. 6**). This enzymatic trait could contribute to the bacterium’s ecological role by enabling it to breakdown complex protein substrates in its environment (Razzaq et al., 2019). Furthermore, the assessment of IAA production through spectrophotometry demonstrated the pink color development upon reaction with the Salkowski reagent, indicating the presence of IAA in the culture supernatant. The concentration of IAA produced by *Luteibacter* strain PPL193^T^ was quantified to be 30.68 ± 0.033 µg/mL (**Fig. 7**). Genome scanning also revealed the presence of IaaH gene in all four strains of the novel species, which is essential for IAA biosynthesis. The IaaH genes among these strains exhibit over 99% sequence identity to each other. IAA, a well-known plant growth-promoting phytohormone, suggests the potential of this novel species to enhance plant growth and development, thus establishing its significance in agricultural applications.

**Fig. 6:**
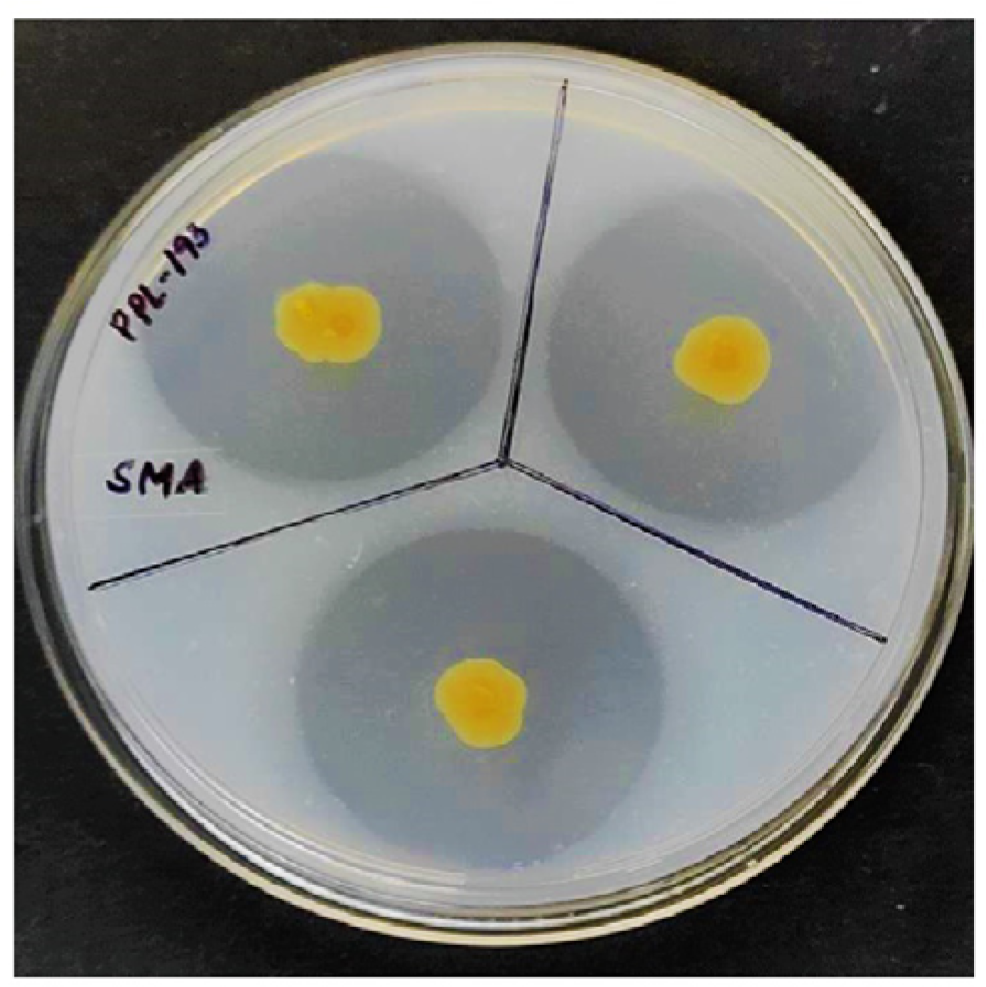
Protease Production by PPL193^T^. Protease activity was assessed using a 1% Skimmed Milk Agar (SMA) medium. The image shows a representative plate with the isolate PPL193^T^ after 10 days of incubation at 28°C. The radius of the clear zone surrounding the colony is 1.36 ± 0.057 cm, indicating the protease-producing capability of PPL193^T^.

**Fig. 7:**
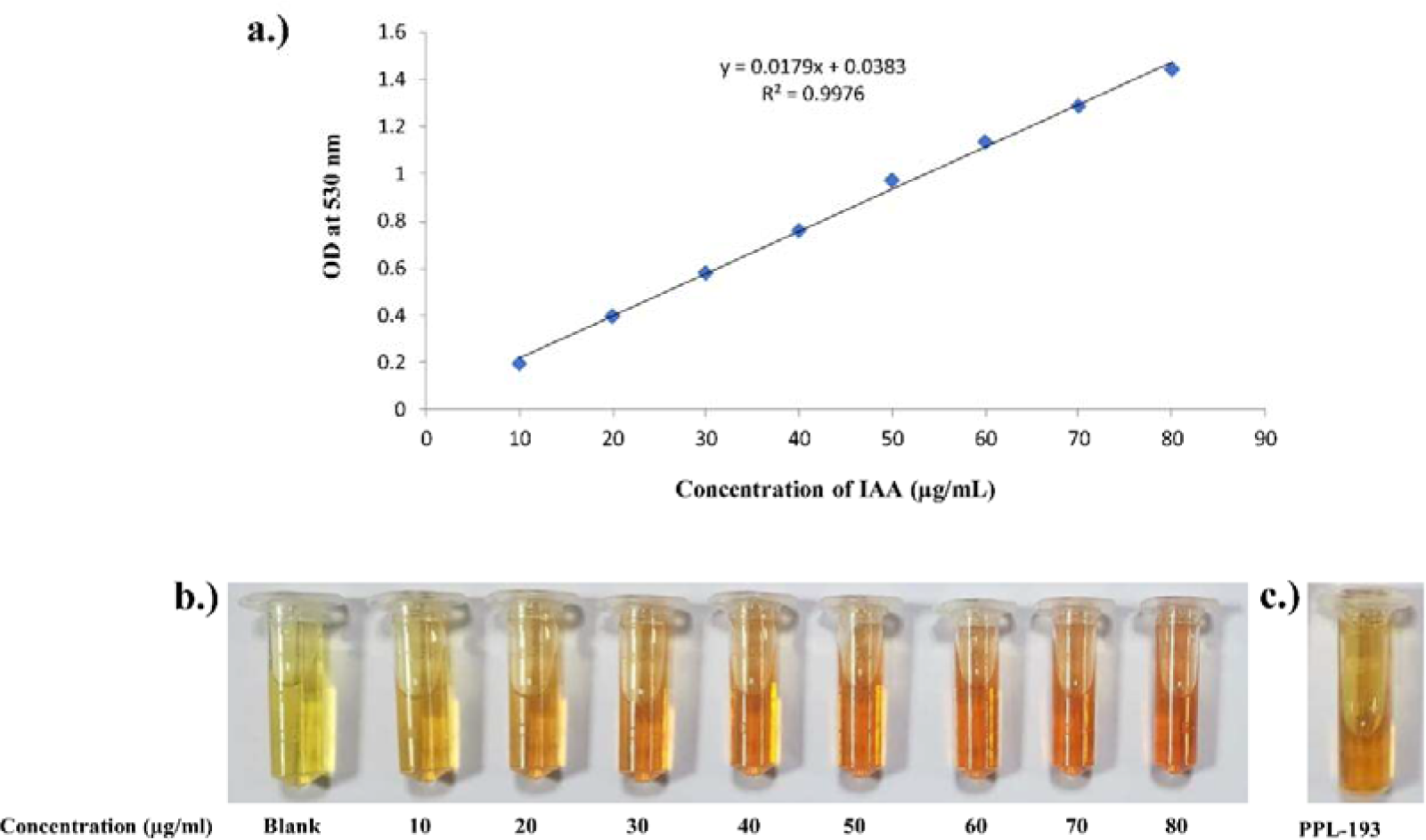
Indole-3-Acetic Acid (IAA) production by PPL193^T^. IAA production was quantified using spectrophotometry after 7 days of incubation in Nutrient Broth (NB) supplemented with 0.2% L-tryptophan at 28°C and 125 rpm. **a.)** A standard curve using solutions with known concentrations of IAA. **b.)** Representative image of solutions with known concentrations of IAA producing pink color. **c.)** Displays the development of a pink color, indicating IAA production by PPL193^T^. The concentration of IAA produced by PPL193^T^ was quantified at 30.68 ± 0.033 µg/ml.

### Assessment of pathogenicity and co-inoculation effects of *Luteibacter* strains PPL193^T^ and PPL552 in rice plants

*Luteibacter* strains PPL193^T^ and PPL552 were derived from healthy rice seeds, prompting an investigation into their pathogenicity through *in-planta* studies via leaf-clip inoculation. As a positive control, *X.oryzae* pv. o*ryzae* (Xoo) strain BXO1, known for causing bacterial blight disease in rice, was used (Bansal et al., 2021). Additionally, plant leaves inoculated solely with MQ water served as a negative control. After 15-days of the infection period, disease symptom/lesion length was measured on the inoculated leaves. As expected, PPL193^T^ and PPL552 isolates did not induce any lesions or disease symptoms on rice leaves, while leaves inoculated with BXO1 showed lesions, with an average length of 15 cm, evident through yellowing and wilting of rice leaves **(Fig. 8a)**. These results indicate that PPL193^T^ and PPL552 isolates are non-pathogenic and natural residents of the host plant.

**Fig. 8:**
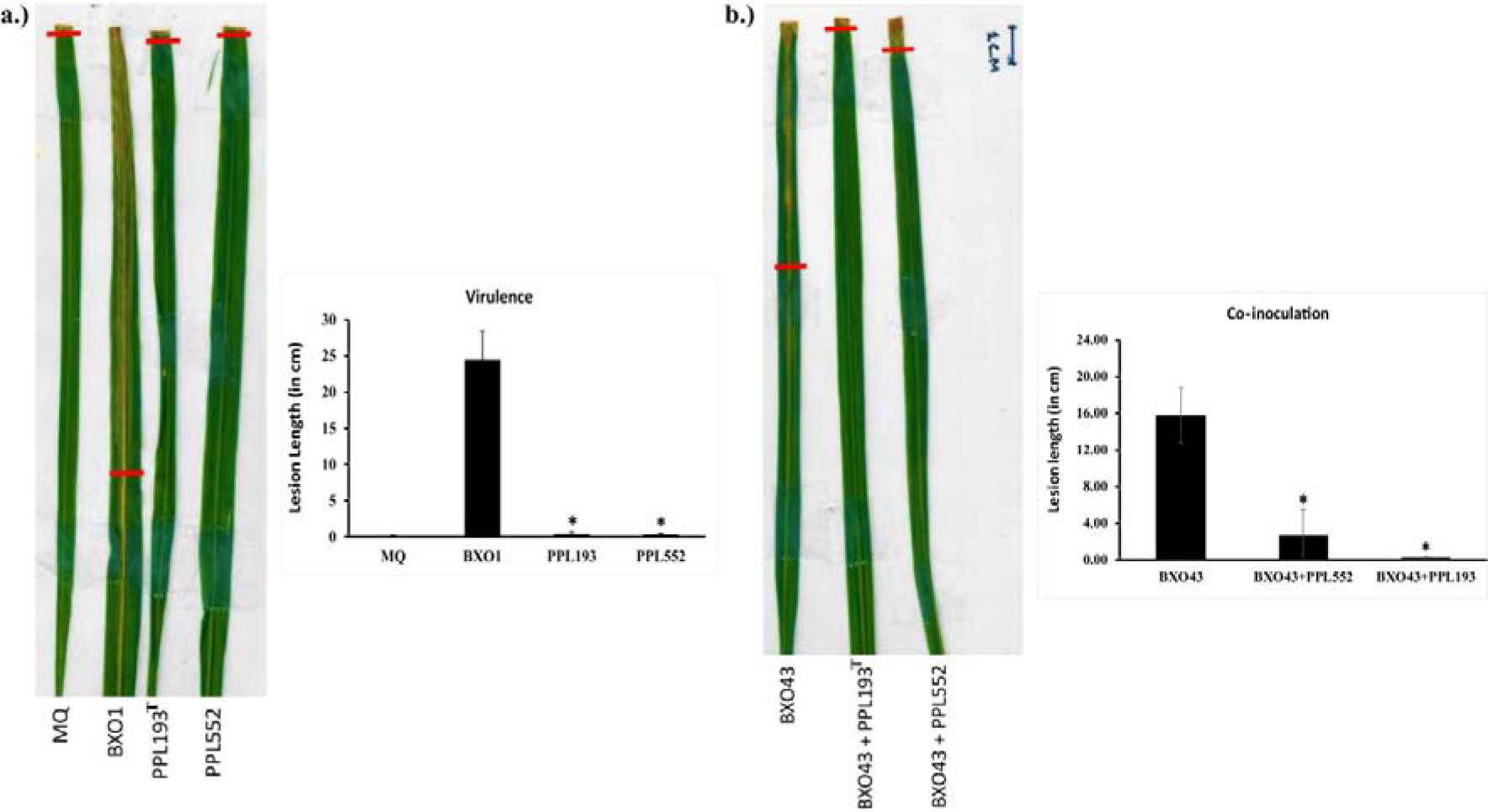
Investigating pathogenicity and evaluating biocontrol potential of PPL193^T^ and PPL552 Strains. The bacterial strains were inoculated on rice leaves following the protocol outlined in the materials and methods section. Figure 8a displays a composite image of rice leaves that were subjected to infection alongside a bar graph illustrating the average lesion length indicating virulence. This comparison includes PPL193^T^ and PPL552 strains against the wild-type BXO1 strain of Xoo. In Figure 8b, a composite image of co-inoculated rice leaves is presented, accompanied by a bar graph representing the average lesion length after the co-inoculation involving the wild-type Xoo strain BXO43 with PPL193^T^ and PPL552. The error bars depict standard deviations calculated from a minimum of 8 leaves. Column labels marked with an asterisk (*) denote significant variations in lesion length, as determined through unpaired two-tailed Student’s t-test (P-value < 0.05).

As these *Luteibacter* strains might be co-evolving with the pathogenic rice strains, we explored the possibility that these strains may provide protection against virulence-proficient pathogenic bacteria and thus benefit the host plant. To delve into this further, we conducted co-inoculation experiments by combining suspensions of *Luteibacter* strains and pathogenic strain of Xoo, BXO43 (rif^R^ derivative of BXO1) (Ray et al., 2000). To our surprise, there was the absence of virulence lesions in the leaves after co-inoculation (**Fig. 8b**), suggesting that the novel *Luteibacter* strains inhibit Xoo growth *in-planta*. This observation adds to our previous findings on the bio-protection property observed in *X. sontii* and *X. indica* (NPX strains) isolated from the healthy rice seeds in our lab (Bansal et al., 2021; Rana et al., 2022).

### Diverse secondary metabolite biosynthetic gene clusters in novel *Luteibacter* species

To obtain further insights into the metabolite potential related to plant growth and protection properties, we scanned the genome for putative secondary metabolite biosynthetic gene clusters (BGCs) using antiSMASH application (Blin et al., 2021). The application of antiSMASH analysis to the genomes of the novel *Luteibacter* species revealed an array of putative secondary metabolite BGCs, encompassing diverse classes such asribosomally synthesized and post-translationally modified (RiPP-like) peptides, non-ribosomal peptide synthetase (NRPS) peptides, polyketide synthases (PKS) compounds, lanthipeptides, arylpolyenes, terpenes, and siderophores (**Table 4**). These identified BGCs hold significant implications for both the ecological fitness and agricultural relevance of the novel species. The representative BGCs for NRPS and lanthipeptide in PPL193^T^ are shown in **Fig. 9**. The distribution of secondary metabolites among the genus *Luteibacter* is shown in **Fig. 10**.

**Fig. 9:**
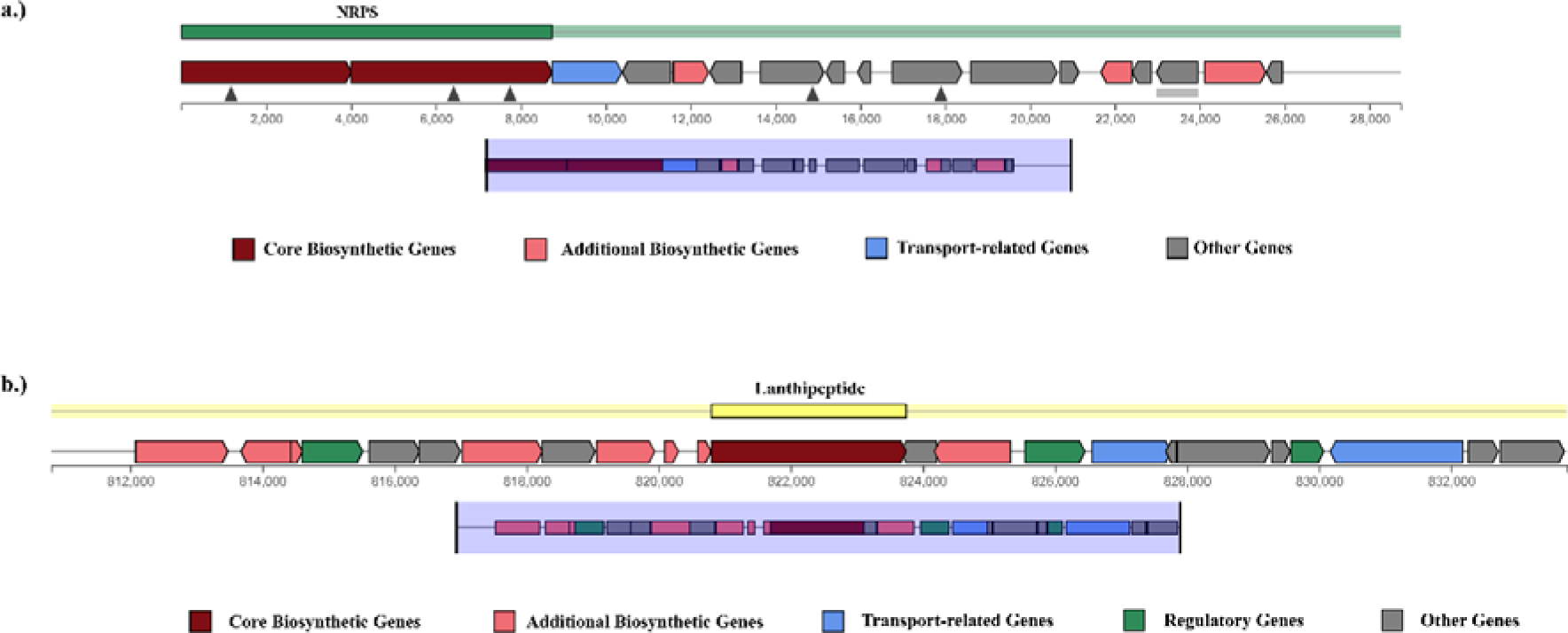
Representative BGCs for NRPS and lanthipeptide within the genome of the novel bacterial species PPL193^T^, providing a snapshot of the NRPS and lanthipeptide biosynthetic potential in PPL193^T^. Highlighted in red are the core biosynthetic genes, while additional biosynthetic genes are marked in pink, transport-related genes in blue, and other associated genes in grey.

**Fig. 10:**
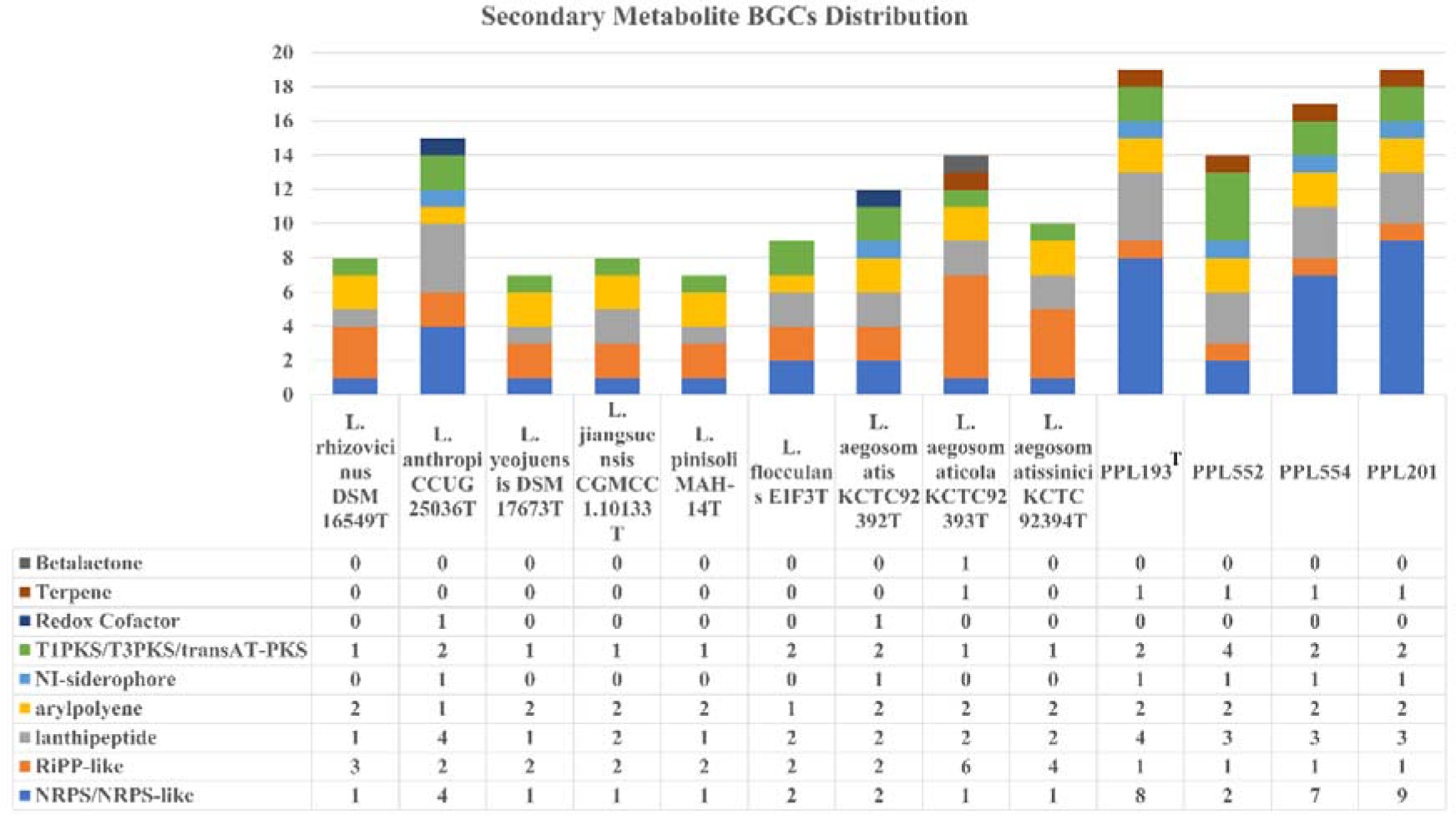
Genomic distribution of secondary metabolite BGCs across the genus *Luteibacter,* highlighting the distribution of distinct BGC classes (each represented by a unique color), including notable classes like NRPS and lanthipeptide, for insights into secondary metabolite diversity. Notably, PPL193^T^and its other constituent members exhibit a higher abundance of secondary metabolite BGCs compared to other members of the genus *Luteibacter*.

**Table 4.** AntiSMASH v7.0.1 analysis of PPL193^T^ shows the presence of secondary metabolites, their type, gene cluster position, size, and predicted product.

| Gene Cluster | Contig | Type | Gene cluster position<br>(Coordinates, From: To) | Size (kb) | Predicted Product (Similarity) |
| --- | --- | --- | --- | --- | --- |
| Region 1.1 | NODE_1_length_1166098_cov_405.353852 | Arylpolyene | 654638-703307 | 48.67 | APE Ec (26%) |
| Region 1.2 |  | Terpene | 735567-756403 | 20.84 | Unknown |
| Region 1.3 |  | Lanthipeptide class II | 810804-833749 | 22.95 | Unknown |
| Region 2.1 | NODE_2_length_1134670_cov_406.543290 | RiPP-like | 1040167-1051012 | 10.85 | Unknown |
| Region 3.1 | NODE_3_length_569792_cov_416.492811 | NRPS | 1-28735 | 28.73 | Kolossin (100%) |
| Region 3.2 |  | Lanthipeptide class III | 82456-105149 | 22.7 | Unknown |
| Region 4.1 | NODE_4_length_242589_cov_409.840948 | NRPS, T1PKS | 109796-162562 | 52.77 | Unknown |
| Region 5.1 | NODE_5_length_219313_cov_410.128223 | Lanthipeptide class IV | 194719-217406 | 22.69 | Unknown |
| Region 6.1 | NODE_6_length_163214_cov_407.861337 | Arylpolyene | 103643-144845 | 41.2 | Unknown |
| Region 8.1 | NODE_8_length_136779_cov_416.499751 | Lanthipeptide class II | 1-14481 | 14.48 | Unknown |
| Region 8.2 |  | TransAT-PKS, NRPS, NI-siderophore | 17996-95701 | 77.71 | Xanthoferrin (100%) |
| Region 9.1 | NODE_9_length_101104_cov_429.921110 | NRPS | 75807-101104 | 25.3 | Unknown |
| Region 14.1 | NODE_14_length_7700_cov_471.257248 | NRPS | 1-7700 | 7.7 | Unknown |
| Region 15.1 | NODE_15_length_5980_cov_513.643063 | NRPS | 1-5980 | 5.98 | Unknown |
| Region 17.1 | NODE_17_length_4841_cov_777.042821 | NRPS | 1-4841 | 4.84 | Rhizomide A/ Rhizomide B/ Rhizomide C (100%) |
| Region 18.1 | NODE_18_length_3454_cov_553.352384 | NRPS | 1-3454 | 3.45 | Nunapeptin/ Nunamycin (35%) |

In particular, the presence of RiPP-like, NRPS, and T1PKS clusters suggests the potential biosynthesis of bioactive molecules with diverse antimicrobial properties (Maiti and Mandal, 2022). These metabolites could play a pivotal role in mediating interactions between the novel *Luteibacter* species and its environment, including potential plant pathogens. Furthermore, the identification of lanthipeptide clusters points to the production of ribosomally synthesized peptides (Alkhalili and Canbäck, 2018), which often exhibit antibacterial activities, reinforcing the plant protection traits observed.

Moreover, the presence of terpene and arylpolyene clusters suggests the capacity for volatile compound production and epiphytic survival, respectively, which may contribute to plant growth promotion through enhancing nutrient uptake and stimulating plant defences (He et al., 2020; Singh and Sharma, 2015). Notably, the siderophore BGCs indicate the ability of the novel *Luteibacter* species to facilitate iron acquisition, a crucial factor in plant health and growth (Wang et al., 2022). Finally, the transAT-PKS clusters hint at the potential production of complex polyketides with diverse biological activities (Almeida et al., 2022), which can further contribute to both plant protection and growth promotion.

The concurrence of these secondary metabolite biosynthetic gene clusters with the observed plant protection and growth promotion traits underscores the multifaceted role of the novel *Luteibacter* species isolated from the rice seeds in plant-microbe interactions. Further investigation of these BGCs and the metabolites they encode promises to illuminate the precise mechanisms underlying the observed phenotypes and open avenues for innovative biocontrol strategies.

### Pangenomic insights and functional characterization of distinct genes associated with PPL193^T^

In this study, we analyzed the pangenome of PPL193^T^, a novel strain belonging to the *Luteibacter* genus. Our findings showed that the pangenome of PPL193^T^ with other type strains of *Luteibacter* spp. contains 28,582 genes, out of which 652 genes were found to be core genes, and 2,274 genes were unique to PPL193^T^ (**Fig. 11a**). These unique genes were functionally annotated and classified into COG categories, providing insights into the potential functions and characteristics of PPL193^T^. Interestingly, we found that approximately 27.3% of the unique genes of PPL193^T^ could not be assigned to any of the COG classes and were classified under the unknown function(s) category (**Fig.11b**). These genes encoded hypothetical proteins or DUF domain-containing proteins, indicating their potential roles in unknown metabolic pathways or functions. On the other hand, the remaining unique genes were found to be involved in various transport and metabolic processes, such as amino acids, nucleotides, inorganic ions, lipids, carbohydrates, and coenzymes. Moreover, the COG categories of energy production and conversion, transcription (K), signal transduction mechanisms (T), and cell wall/membrane/envelope biogenesis (M) were found to contain a significant proportion of unique genes of PPL193^T^, approximately 6-9% each. Additionally, we compared the COG classification of unique genes of PPL193^T^ with KCTC92392^T^, another *Luteibacter* strain, and found that approximately 27-29% of the unique genes were not assigned to any COG class or were classified under the unknown function (S) category in both strains. However, PPL193^T^ carried more unique genes involved in nucleotide transport and metabolism (F), replication, recombination, and repair (L), cell wall/membrane/envelope biogenesis (M), and cell motility (N) categories, as compared to KCTC92392^T^ **(Fig.11c)**.

**Fig. 11:**
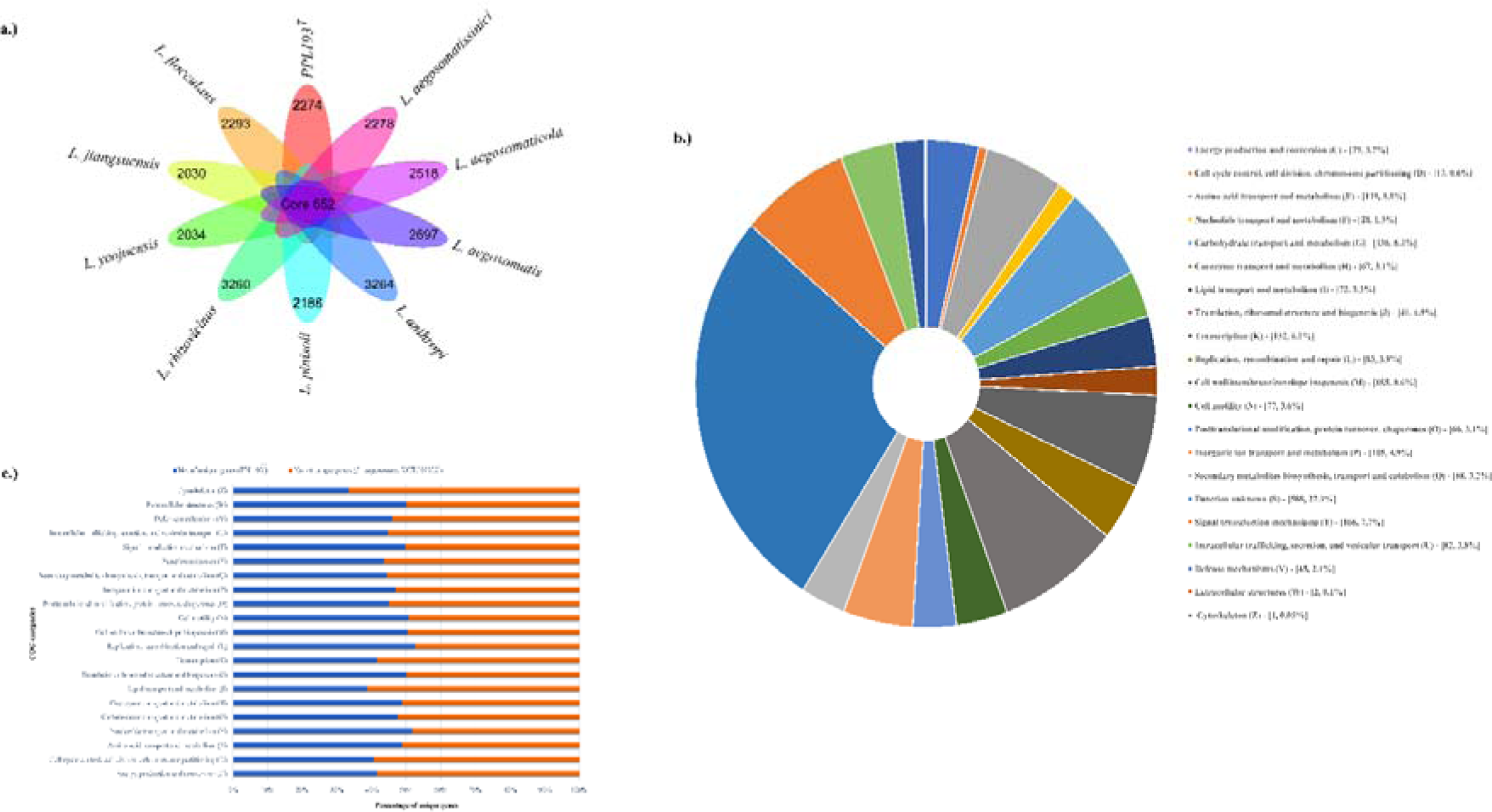
Unveiling the genomic landscape through pan-genome analysis of PPL193^T^, the type strain. **11a.**Visualizing the core and unique genes of the novel isolate PPL193^T^ alongside other *Luteibacter* spp. **11b.** Illustrating the distribution of PPL193^T^’s unique genes across various COG categories. **11c.** A bar graph revealing the functional annotation of PPL193^T^’s unique genes and *Luteibacter aegosomatis* KCTC 92392^T^, categorized into COG categories.

## Conclusion

In conclusion, we report the discovery and characterization of novel yellow-pigmented *Luteibacter* isolates from healthy rice seeds. The comprehensive characterization of the isolated strain PPL193^T^ using phenotypic, biochemical, 16S rRNA gene sequence investigation, along with genome-based taxonomy, phylogenetic, and comparative analysis, revealed that the strain PPL193^T^represents a new species within the genus *Luteibacter* along with PPL201, PPL552, and PPL554 as its members. These findings provide important insights into the unique genomic characteristics and potential metabolic pathways of PPL193^T^, which could contribute to the understanding of its adaptation and evolution from a novel ecological niche of importance to seed health. Moreover, these results could have implications for the development of consortia for plant protection and exploitation of seed microbiome as revealed by plant growth and protection properties of the isolates from the new species. Interestingly, the finding of the production of xanthomonadin pigment and also genomic loci that is under variation in the genus indicates ongoing evolution towards adaption to diverse niches. Further genetic and chemical characterization of the pigment gene cluster will aid in our understanding of the evolution of this locus in *Xanthomonas* and its taxonomic/ecological relatives. These findings expand our understanding of the diversity and ecology of bacterial communities associated with rice seeds. This study also contributes to the growing body of literature on taxonogenomics and provides valuable insights into the genomic characteristics of these novel *Luteibacter* isolates.

### Description of *Luteibacter sahnii* sp. nov

*Luteibacter sahnii* (sah’ni.i. N.L. masc. gen. n. sahnii) has been named in honour of Girish Sahni, a renowned microbiologist, science leader and administrator from India with specialization in protein engineering, molecular biology, and biotechnology.

Isolates of this species form convex, light-yellow, and mucoid colonies following 48 hours of incubation at 28°C on the NA medium. Cells of this Gram-negative, aerobic, and rod-shaped bacterium can utilize D-maltose, D-trehalose, D-cellobiose, gentiobiose, sucrose, D-turanose, α-D-lactose, N-Acetyl-D-Glucosamine, α-D-glucose, D-mannose, L-Fucose as the carbon sources. The species was delineated from other reported species through phylo-taxonogenomic characterization. Growth was observed at pH 5.0 and 6.0, 4% NaCl, and 1% sodium lactate. The genome size and G+C content of the type strain are 4.21 Mb and 67%, respectively. The isolates from the species are non-pathogenic to rice plants with plant growth and protection properties. The type strain of this species is PPL193^T^= MTCC 13290^T^=ICMP 24807^T^=CFBP 9144^T^, and PPL201, PPL552, and PPL554 are other constituent members of the novel species.

